# Genome-wide association study and functional validation implicates JADE1 in tauopathy

**DOI:** 10.1101/2021.06.30.450599

**Authors:** Kurt Farrell, SoongHo Kim, Natalia Han, Megan A. Iida, Elias Gonzalez, Marcos Otero-Garcia, Jamie Walker, Tim Richardson, Alan E. Renton, Shea J. Andrews, Brian Fulton-Howard, Jack Humphrey, Ricardo A. Vialle, Kathryn R. Bowles, Kristen Whitney, Diana K. Dangoor, Edoardo Marcora, Marco M. Hefti, Alicia Casella, Cheick Sissoko, Manav Kapoor, Gloriia Novikova, Evan Udine, Garrett Wong, Weijing Tang, Tushar Bhangale, Julie Hunkapiller, Gai Ayalon, Rob Graham, Jonathan D. Cherry, Etty Cortes, Valeriy Borukov, Ann C. McKee, Thor D. Stein, Jean-Paul Vonsattel, Andy F. Teich, Marla Gearing, Jonathan Glass, Juan C. Troncoso, Matthew P. Frosch, Bradley T. Hyman, Dennis W. Dickson, Melissa E. Murray, Johannes Attems, Margaret E. Flanagan, Qinwen Mao, M-Marsel Mesulam, Sandra Weintraub, Randy Woltjer, Thao Pham, Julia Kofler, Julie A. Schneider, Lei Yu, Dushyant P. Purohit, Vahram Haroutunian, Patrick R. Hof, Sam Gandy, Mary Sano, Thomas G. Beach, Wayne Poon, Claudia Kawas, María Corrada, Robert A. Rissman, Jeff Metcalf, Sara Shuldberg, Bahar Salehi, Peter T. Nelson, John Q. Trojanowski, Edward B. Lee, David A. Wolk, Corey T. McMillan, Dirk C. Keene, Thomas J. Montine, Gabor G. Kovacs, Mirjam I. Lutz, Peter Fischer, Richard J. Perrin, Nigel Cairns, Erin E. Franklin, Herbert T. Cohen, Maria Inmaculada Cobos Sillero, Bess Frost, Towfique Raj, Alison Goate, Charles L. White, John F. Crary

## Abstract

Primary age-related tauopathy (PART) is a neurodegenerative tauopathy with features distinct from but also overlapping with Alzheimer disease (AD). While both exhibit Alzheimer-type temporal lobe neurofibrillary degeneration alongside amnestic cognitive impairment, PART develops independently of amyloid-β (Aβ) deposition in plaques. The pathogenesis of PART is unknown, but evidence suggests it is associated with genes that promote tau pathology as well as others that protect from Aβ toxicity. Here, we performed a genetic association study in an autopsy cohort of individuals with PART (*n*=647) using Braak neurofibrillary tangle stage as a quantitative trait adjusting for sex, age, genotyping platform, and principal components. We found significant associations with some candidate loci associated with AD and progressive supranuclear palsy, a primary tauopathy (*SLC24A4, MS4A6A, HS3ST1, MAPT* and *EIF2AK3*). Genome-wide association analysis revealed a novel significant association with a single nucleotide polymorphism on chromosome 4 (rs56405341) in a locus containing three genes, including *JADE1* which was significantly upregulated in tangle-bearing neurons by single-soma RNA-seq. Immunohistochemical studies using antisera targeting JADE1 protein revealed localization within tau aggregates in autopsy brain from tauopathies containing isoforms with four microtubule-binding domain repeats (4R) and mixed 3R/4R, but not with 3R exclusively. Co-immunoprecipitation revealed a direct and specific binding of JADE1 protein to tau containing four (4R) and no N-terminal inserts (0N4R) in post-mortem human PART brain tissue. Finally, knockdown of the *Drosophila* JADE1 homolog rhinoceros (rno) enhanced tau-induced toxicity and apoptosis *in vivo* in a humanized 0N4R mutant tau knock-in model as quantified by rough eye phenotype and terminal deoxynucleotidyl transferase dUTP nick end-labeling (TUNEL) in the fly brain. Together, these findings indicate that PART has a genetic architecture that partially overlaps with AD and other tauopathies and suggests a novel role for JADE1 as a mediator of neurofibrillary degeneration.

## Introduction

Primary age-related tauopathy (PART) is nearly ubiquitously observed with varying degrees of severity in the brains of aged individuals characterized by the presence of neurofibrillary tangles (NFT) composed of abnormal aggregates of tau protein. These NFTs are regionally, morphologically, biochemically and ultrastructurally identical to those in early to moderate stage Alzheimer disease (AD), yet develops in the absence of Aβ plaques ^1^. A neuropathological diagnosis of PART is separate from a clinical diagnosis of cognitive impairment; hence individuals with PART can be normal, mildly cognitively impaired (MCI) or have dementia ^2,3^. Most individuals with PART remain cognitively normal, however some develop amnestic cognitive changes ^4-7^. The similarities between PART and AD allows for a unique opportunity to focus specifically on mechanisms of tau-mediated AD-type neurodegeneration. The neuropathological diagnosis of PART is complicated by accompanying age-related comorbid dementing pathologies, therefore discovering unique molecular drivers is challenging antemortem ^8-13^. Given the similarities between PART and AD NFTs, PART as a diagnostic construct would have greater value if it were shown to arise independently ^14,15^. Alternatively, PART might be a component of the AD spectrum, and Aβ pathology might have eventually developed in such individuals had they lived longer ^15,16^. Thus, the question remains as to what extent a neuropathological diagnosis of PART diverges from or has similar risk factors to AD and other dementias ^17,18^.

Much of the mechanistic knowledge surrounding tauopathy stems from genetic studies ^19^. Autosomal dominant mutations in the microtubule-associated protein tau gene (*MAPT*) in coding regions can interfere with microtubule binding or promote transition to toxic forms. Also, mutations in splice sites disrupt alternative pre-mRNA splicing of the tau mRNA transcript, modifying the ratio of three repeat (3R) and four repeat (4R) tau isoforms leading to downstream pathological sequelae ^20^. Additionally, alternative splicing of the N-terminal exons may also play a role in modulating toxic tau ^21,22^. However, *MAPT* mutations are rare, whereas PART occurs sporadically and nearly ubiquitously with advanced age. Thus, PART provides a unique opportunity to understand common genetic drivers of tauopathy and their association with abnormalities in tau proteostasis and isoform expression. Prior research focusing on common genetic variation has identified two distinct haplotypes in the *MAPT* 17q21.31 locus defined by a large ∼900kb inversion region that gives rise to two major haplotypes, H1 and H2. The more common H1 haplotype has been associated with increased risk for PART and several other sporadic tauopathies including *APOE* L4-negative AD, progressive supranuclear palsy, corticobasal degeneration, and even Parkinson disease which is not classically considered a tauopathy ^23-28^. Research focusing on the genetics of PART have consistently failed to show an association with the *APOE* ε4 allele, the strongest risk locus in AD ^28-31^. Outside of PART, one of the largest AD GWAS has identified 29 risk loci implicating 215 potential causative genes ^32^. However, it has been suggested that the overwhelmingly strong signal on the *APOE* locus could mask associations independently related to tauopathy ^23^. Another study examining candidate genes in aging cohorts confirmed the lack of the association between *APOE* ε4 and PART, as well as a decreased frequency of AD risk alleles in the *BIN1, PTK2B*, and *CR1* loci ^33^. Taken together, these data suggest an unexplored genetic risk driving tauopathy that might be revealed by conducting genome-wide association studies in PART.

While PART represents a compelling opportunity to focus on amyloid-independent mechanisms of neurofibrillary degeneration, assembling an autopsy cohort large enough to assesses its genetics on a genome-wide scale has not yet been attempted. In collaboration with twenty-one domestic and international brain biorepositories, we performed the first genome-wide association study in PART and compared our findings to known tauopathy risk loci. We then localized expression of candidate genes in our strongest risk locus (chromosome 4q28.2) using single cell RNA-sequencing and immunohistochemistry. Finally, we validated our findings biochemically in human postmortem brain and functionally using an *in vivo* Drosophila model. The work presented here not only furthers the investigation of the genetics of PART, but also suggests a novel role for *JADE1* in tauopathy.

## Materials and methods

### Cohort

Fresh-frozen brain tissue was obtained from the contributing centers (Supplementary Table 1). All tissue was used in accordance with the relevant guidelines and regulations of the respective institutions. The neuropathological assessments were performed at the respective centers using standardized criteria outlined by the National Institutes on Aging-Alzheimer’s Association (NIA-AA) and National Alzheimer Coordinating Center (NACC) which has a high degree of interrater reliability among centers ^34,35^. Inclusion criteria were individuals with normal cognition, mild cognitive impairment (any type) and dementia. Cognitive status was determined either premortem or postmortem by a clinical chart review, mini-mental score, or clinical dementia rating ^36,37^. Both sexes were included and ages ranged from 51 to 108 years at death. Neuropathological inclusion criteria were Braak tangle stage of 0-IV and neuritic amyloid plaque severity CERAD score of 0 ^38,39^. In addition, tissue sections were obtained and reevaluated by the study investigators to confirm the lack of amyloid and degree of PART tau pathology^40^. Clinical exclusion criteria were motor neuron disease, parkinsonism, and frontotemporal dementia. Neuropathological exclusion criteria were other degenerative diseases associated with NFTs (i.e., AD, progressive supranuclear palsy [PSP], corticobasal degeneration [CBD], chronic traumatic encephalopathy [CTE], frontotemporal lobar degeneration-tau [FTLD-tau], Pick disease, Guam amyotrophic lateral sclerosis/parkinsonism-dementia, subacute sclerosing panencephalitis, globular glial tauopathy). Individuals with aging-related tau astrogliopathy (ARTAG) were not excluded ^41^.

### Genotyping

High-throughput isolation of DNA was performed using the MagMAX DNA Multi-Sample Ultra 2.0 Kit on a KingFisher Flex robotic DNA isolation system (Thermofisher, Waltham, MA). Fresh frozen cortical brain tissue (20-40 mg) was placed into a deep-well plate and treated with 480 µl of Proteinase K mix (Proteinase K, phosphate buffered saline [pH 7.4], Binding Enhancer) and incubated overnight at 65°C at 800 rpm on a shaking plate. Genomic DNA was isolated and purified using magnetic particles. DNA quality control was performed using a nanodrop spectrophotometer (concentration > 50ng/µl, 260/280 ratio 1.7-2.2). Genotyping was performed using single nucleotide polymorphism (SNP) microarrays (Infinium Global Screening Array v2.4. or Infinium OmniExpress-24; Illumina, San Diego, CA). Raw genotype files were converted to PLINK-compatible files using GenomeStudio software (Illumina, San Diego, CA).

### Genetic analysis

PLINK v1.9 was used to perform quality control ^42^. SNP exclusion criteria included minor allele frequency <1%, genotyping call-rate filter less then 95%, and Hardy-Weinberg threshold of 1×10^−6 43^. Individuals with discordant sex, non-European ancestry, genotyping failure of >5%, or relatedness of >0.1 were excluded. A principal component analysis (PCA) was performed to identify population substructure using EIGENSTRAT and the 1000 genomes reference panel ^44,45^. Samples were excluded if they were five standard deviations away from the European population cluster (Supplementary Fig. 4). PCA was performed again on the genomic data with non-Europeans excluded to generate new PCs, of which the first four were used as covariates in the regression model. All data was imputed on the University of Michigan server using minimac3 and HRC reference panel ^46,47^. Imputed variants with MAF <0.01 and a dosage R^2^ <0.7 were excluded. Downstream analyses were based on the most likely genotype. A quantitative trait association test was run on 647 PART cases vs. Gaussian-normalized Braak stage using conditional linear regression and age, sex, principal component 1-4 and SNP chip array as covariates. The analysis was run separately on each genotyping array and a meta-analysis was performed using METAL ^48^. Regional genome-wide association plots were created with LocusZoom, other plots were created using R ^49^.

### Single-cell mRNA profiling in tangle-containing neurons

Identification of differentially expressed genes in single neuronal somata with and without NFTs was performed by analyzing a transcriptomic dataset of isolated neurons from post-mortem human brain from individuals with and without AD reported by Otero-Garcia *et. al*. 2021 ^50^. This dataset consists of single neuronal soma transcriptomes from Brodmann area 9 subjected to fluorescence-activated cell sorting (FACS) using p-tau (AT8) and MAP2 antisera to differentiate single NFT-positive and NFT-negative cells.

### Immunohistochemistry

Formalin-fixed paraffin-embedded tissue sections (5 μm) on charged slides were baked at 70°C and immunohistochemistry (IHC) was performed on a Ventana Benchmark XT automatic stainer (Rouche, Tucson, AZ). Antigen retrieval was done using citric acid buffer (CC1) for 1 hr followed by primary antibody incubation for approximately 40 min. A secondary antibody, 3,3’-diaminobenzidine (DAB) was then applied. For slides that were doubled-labeled DAB and alkaline phosphatase were used for visualization. Slides were stained with antibodies to JADE1 (1:100, Proteintech, Rosemont, IL) and hyperphosphorylated tau (p-tau, AT8, 1:1000, Invitrogen, Waltham, MA). To ensure specificity of the JADE1 antisera, a peptide competition was performed using a blocking peptide. The antisera and paired peptide were pre-incubated for 24 hr. Whole slide images (WSI) were visualized and scanned using an Aperio CS2 digital slide scanner (Leica Biosystems, Wetzlar Germany). In addition to PART cases, neuropathologically confirmed cases of AD, PSP, CTE, CBD, argyrophilic grain disease (AGD) and Pick disease (PiD) (*n*=3 for each, Supplemental table 6) were examined for convergent or divergent staining patterns.

### Biochemical analysis

Western blots were performed using homogenized fresh-frozen brain tissue. Samples were placed in a micro-tube homogenizer (SP Bel-Art, Wayne, NJ) in 10 volumes (wt/vol) of ice-cold Pierce RIPA buffer (Thermo Fisher Scientific, Waltham, MA) containing Halt protease and phosphatase inhibitor cocktail (Thermo Fisher Scientific), and incubated on ice for 30 min. For each sample, 20 μg of proteins were boiled in 1x Laemmli sample buffer (Bio-Rad, Hercules, CA) for 5 min, run on 10% Criterion TGX Precast Gels (Bio-Rad, Hercules, CA), blotted to nitrocellulose membranes, and stained with JADE1 antisera (1:2000). Horseradish peroxidase-labeled secondary anti-rabbit antisera (1:20,000; Vector Labs, Burlingame, CA) was detected by SuperSignal West Pico PLUS Chemiluminescent Substrate or Pierce ECL Western Blotting Substrate (Thermo Fisher Scientific). To quantify and standardize protein levels, GAPDH was detected with GAPDH antisera (6C5, 1:20,000; Abcam, Cambridge, MA) and total protein was detected with Amido Black (Sigma-Aldrich, St. Louis, MO) as previously described ^51^. Chemiluminescence was measured in a ChemiDoc Imaging System (Bio-Rad), and relative optical densities were determined by using AlphaEaseFC software, version 4.0.1 (Alpha Innotech, San Jose, CA), normalized to GAPDH and total protein loaded.

Co-immunoprecipitation (IP) assays were performed using fresh-frozen brain tissue was homogenized in a glass-Teflon homogenizer at 500 rpm in 10 volumes (wt/vol) of ice-cold lysis buffer containing 50 mM Tris, pH 7.8, 0.5 % NP40, 150 mM NaCl, 1 mM EDTA, and Halt protease and phosphatase inhibitor cocktail (Thermo Fisher Scientific). Samples were incubated on ice for 30 min, centrifuged at 1000 x g for 10 min and the supernatant was collected as an input and used for immunoprecipitation. In a microcentrifuge tube, 70 ul of supernatant, Lysis buffer (930 µl) and 2 µg of either JADE1 antisera or anti-0N tau antisera (EPR21726, Abcam, Cambridge, MA) were combined and incubated overnight at 4 °C. Two controls were also set up, one without the antisera and the other with 2 µg of IgG isotype control antisera, either normal rabbit IgG (PeproTech, Rocky Hill, NJ) or Mouse IgG1 kappa, (clone: P3.6.2.8.1, Thermo Fisher Scientific). Twenty µl of Pierce Protein A/G Agarose beads (Thermo Fisher Scientific) was added to each reaction, and the mixture was incubated for 1 hr at 4 °C. Agarose beads were pelleted at 1000 x g for 5 min at 4 °C, supernatant was removed, 1 ml of ice-cold lysis buffer was added, and pellet was washed by inverting tube several times. Beads were washed 4 times, each time repeating the centrifugation step above. After the final wash, pelleted beads were resuspended in 40 µl of 1x Laemmli sample buffer (Bio-Rad, Hercules, CA) and boiled for 5 min. The samples were then centrifuged to pellet the agarose beads followed by SDS-PAGE analysis of the supernatant. Fifteen µl of samples for JADE1 detection and 5 µl for tau with tau isoform ladders (rPeptide, Watkinsville, GA) were run on 10% PROTEAN TGX Precast Gels (Bio-Rad, Hercules, CA), blotted to nitrocellulose membranes, and stained with JADE1 antisera (1:2000), total tau antisera (HT7, 1:3000; Thermo Fisher Scientific), three microtubule repeat domain tau antisera (3R, 8E6/C11, 1:2000; MilliporeSigma, St. Louis, MO), four microtubule repeat domain tau antisera (4R, 1:2000; Cosmo Bio, Carlsbad, CA), pThr231 tau antisera (RZ3, 1:200; a gift from Dr. Peter Davies), pThr181 tau antisera (PHF1, 1:500; a gift from Dr. Peter Davies), pSer202 tau antisera (CP13, 1:500; a gift from Dr. Peter Davies), pSer202/pThr305 tau antisera (AT8, 1:1000; Thermo Fisher Scientific), pSer214 tau antisera (S214; 44-742G, 1:1000; Thermo Fisher Scientific, Waltham, MA), or pSer409 tau antisera (PG5, 1:200; a gift from Dr. Peter Davies). Horseradish peroxidase-labeled conformation-sensitive secondary anti-mouse IgG for IP or anti-rabbit VeriBlot for IP Detection antibody (both at 1:20000; Abcam, Cambridge, MA) was detected by SuperSignal West Femto Maximum Sensitivity substrate (Thermo Fisher Scientific).

A dephosphorylation assay was performed using fresh-frozen brain tissue homogenized with a glass-Teflon homogenizer at 500 rpm in 10 volumes (wt/vol) of ice-cold lysis buffer containing 50 mM Tris, pH 7.8, 0.5 % NP40, 150 mM NaCl, and Halt protease inhibitor cocktail (Thermo Fisher Scientific, Waltham, MA). Samples were incubated on ice for 30 min, centrifuged at 1000 x g for 10 min, and supernatant was collected. Reaction mixtures (51 µul) consisted of 39 µl of supernatant, 1 µl of protease inhibitor cocktail, 5 µl of 10X NEBuffer for protein metallophosphatases, 5 µl of 10 mM MnCl_2_, 1 µl of lambda protein phosphatase (New England BioLabs, Ipswich, MA). Each mixture was incubated at 30 °C for either 1, 2, 3, or 4 h. Additional 1 µl of lambda protein phosphatase and 1 µl of protease inhibitor cocktail were added in each mixture every hour.

### Proximity ligation assay

A proximity ligation assay was performed on formalin-fixed paraffin embedded 5 μm-thick hippocampal sections mounted on charged slides using a Duolink kit (MilliporeSigma, St. Louis, MO). Sections were deparaffinized and incubated in sodium citrate buffer (10 mM sodium citrate, 0.05 % Tween 20, pH 6.0) at 95 °C for 20 min, washed in running water, incubated in 0.2 % Tween 20 in PBS at room temperature for 20 min, and washed in PBS 3 times for 5 minutes.. From blocking, assays were performed using the *in situ* red starter kit according to the manufacture’s protocol with JADE1 antisera (1:20) and 0N tau antisera (1:500; BioLegend, San Diego, CA). Two control assays were also performed, one with JADE1 antisera only, and the other with anti-0N tau antisera. All samples were counterstained with 4′,6-diamidino-2-phenylindole (DAPI). Sections were imaged using an Axioview fluorescent microscope (Carl Zeiss, Oberkochen, Germany) and processed using Zen Blue software.

### In vivo Drosophila model

*Drosophila* stocks, crosses, and aging were performed at 25 °C for the duration of the experiment and an equal number of male and female flies were used for each experiment. The GAL4-UAS expression system and the pan-neuronal elav-GAL4 driver were used to control transgenic human tau and rno expression. Analyses were run on four fly groups (Bloomington stock rno^RNAi^ line number 57774): elav-GAL4 driver in the heterozygous state (control, elav-GAL4/+), elav-Gal4 positive plus rno^RNAi^ positive group (control + rno^RNAi^, elav-GAL4/+;UAS-rno^RNAi^/+), elav-Gal4 positive transgenic human UAS-tau^R406W^ tau group (0N4R Tau, elav-GAL4/+;UAS-Tau^R406W^), elav-Gal4 positive human transgenic UAS-tau^R406W^ tau plus rno^RNAi^ positive group (0N4R Tau + rno^RNAi^, elav-GAL4/+;UAS-Tau^R406W^/UAS-rno^RNAi^). An additional rno^RNAi^ lines were used but was did not produce progeny when crossed to tau transgenic *Drosophila*, as often occurs with strong enhancers (Bloomington stock rno^RNAi^ line number 62880). Flies were aged to 10 days, at which point terminal deoxynucleotidyl transferase dUTP nick end-labeling (TUNEL) assay was performed in the fly brain (*n*=6 per genotype) and a blinded assessment of the fly eye phenotype was performed (*n=*16 total). TUNEL was performed on 4 μm formalin-fixed paraffin-embedded fly heads. Quantification of TUNEL-positive nuclei was performed throughout the entire brain using DAB-based detection and bright field microscopy. Fly eye phenotype scoring was performed blindly using light microscopy (*n*=16). The blinded rater semi-quantitively evaluated the eye for four distinct qualities including roughness, size, shape and conical shape on a 1 to 5 scale (five being the most severe phenotype) and an average summary score was calculated.

### Statistical analysis

For GWAS, statistical analysis was performed in PLINK and our genome wide significance value was < 5 × 10^−8^, which is Bonferroni-corrected for all the independent common SNPs across the human genome. Genome wide suggestive significant value was set at < 5 × 10^−6^. All other statistical analyses were performed in R. For non-normally distributed data a Wilcox test was used to test for significance, and an ANOVA was used for normally distributed data.

## Results

We assembled the first cohort of individuals with primary age-related tauopathy (PART) in which all individuals were neuropathologically confirmed to be devoid of any neuritic plaques. Autopsy brain tissue samples (*n*=647) were obtained from 21 brain banks. While each center had performed a comprehensive neurodegeneration workup, we restained and reassessed these cases for PART pathology as a component of our ongoing histological studies^40^. Neurofibrillary tangles (NFTs) were assessed using the Braak staging system which ranged from 0-IV with all stages well represented, as is consistent with PART (Table 1). The average age of death was 83 years old, and the number of male and females in the study was approximately equal. Amongst the assigned Braak stages, stage II had the highest abundance (*n*=189), relatively equal amounts in stages III (*n*=152) and I, (*n*=142), and the lowest in stages 0 (n=71) and IV (*n*=93). Sixty-six percent of the cases were cognitively normal, however amongst stages I-IV, there was an equal number of cognitively impaired individuals. Braak staging performed at each center was not disproportionally skewed (Supplementary Fig. 1a). Lastly, we investigated the effect of age of death on Braak stage with respect to cognitive status and found a positive correlation that does not significantly change when cognitive status is accounted for in the model (Supplementary Fig. 1a,b). In summary, our cohort consists of primarily older individuals, with a range of clinical cognitive symptoms, as well as a broad spectrum of regional tau distribution, demonstrating the diversity of both the clinical and neuropathological features of the condition.

**Table 1.** Subject data stratified of Braak stage

| Braak NFT stage | n | Average age (range) | Male (%) / Female (%) | Normal (%) / MCI (%) / Dementia (%)* |
| --- | --- | --- | --- | --- |
| 0 (71) | 71 | 71.2 (51-104) | 48 (7.5) / 23 (3.5) | 58 (8.9) / 7 (1.0) / 6 (0.9) |
| I (142) | 142 | 78.1 (53-108) | 88 (13.6) / 54 (8.3) | 99 (15.3) / 17 (2.6) / 22 (3.4) |
| II (189) | 189 | 84.1 (51-107) | 98 (15.1) / 91 (14.0) | 126 (19.4) / 40 (6.1) / 21 (3.2) |
| III (152) | 152 | 88.8 (68-105) | 59 (9.1) / 93 (14.8) | 96 (14.8) / 29 (4.4) / 26 (4.0) |
| IV (93) | 93 | 91.2 (67-106) | 29 (4.4) / 64 (9.8) | 50 (7.7) / 22 (3.4) / 21 (3.2) |
| **Total (647)** | **647** | 83.5 (51-108) | 322 (49.7) / 325 (50.2) | 429 (66.3) / 115 (17.7) / 96 (14.8) |

\*7 samples did not have cognitive status available, MCI = mild cognitive impairment

Using this cohort, we ran a quantitative trait association analysis across the entire genome to identify novel genetic risk loci in PART. Using Braak stage as a quantitative trait revealed a genome wide associated signal on chromosome 4q28.2 (Fig. 1a, rs56405341; linear regression β=0.35, standard error=0.06, *p*=4.82 × 10^−8^) and suggestive signals in 14 other loci (Table 2 and Supplementary Table 3). Our model, which adjusted for age, sex, and genotyping platform and principal component 1-4 produced a λ of 1.04 (Supplementary Fig. 2a). The genome wide significant variant (rs56405341) has a min frequency of 0.27. The locusZoom plot indicates our significant SNP is not directly in an intron r allele of any specific gene, but near *C4orf33, SCLT1* and *JADE1* (Figure 1b). We also observed in our regional plot that the lead SNP has a total of 22 other SNPs in Linkage Disequilibrium (r^2^ > 0.8, Supplementary table 4). Further examination of the homozygous and heterozygous alleles using strip chart shows the significant relationship between higher Braak stage and homozygous minor allele carriers (Figure 1 c, AA-AG *p*=0.024, AA-GG *p*=3.3 × 10^−5^, AG-GG *p*=7.2 × 10^−5^). Separate analyses two different genotyping chip validated our findings by showing replication of the signal on each genotyping platform, as well as comparable lambda values (Infinium OmniExpress-24, n=440, β□=□0.27, SE□=□0.05, *p*□=□1.11× 10^−6^, λ = 1.03, Global screening array β□=□0.20, SE□=□0.08, *p*□=□1.42× 10^−2^, λ = 1.01, Supplementary Table 2 and Fig. 5 a-d). The individual summary statistics derived from the separate chip analysis were then combined to run a meta-analysis, and the resulting p-value was similar to value on the combined analysis, as agreement in the direction of effect tested allele (*p*□= 5.61 × well as 10^−8^).

**Fig. 1.**
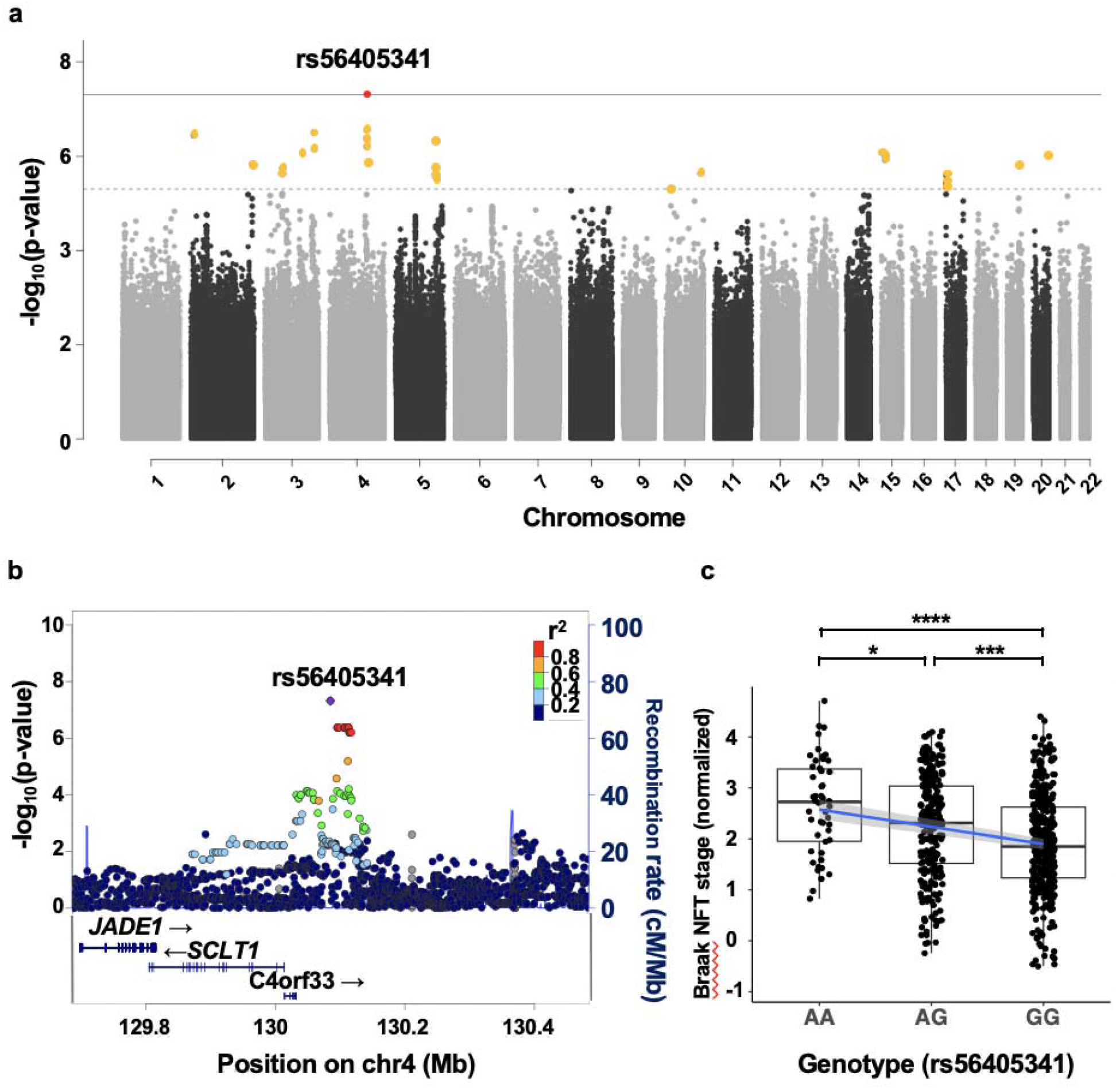
Genome-wide association study (GWAS) in primary age-related tauopathy. (**a**) A quantitative trait GWAS was performed using normalized Braak neurofibrillary tangle stage with age, sex, principle components (PCs) and genotyping SNP array as covariates (*n*=647). The threshold for genome-wide significance (*p*<5 × 10^−8^) is indicated by the solid grey line; the suggestive line (*p*<5 × 10^−6^) is indicated by the dotted line. (**b**) A LocusZoom plot shows a strong signal with multiple SNPs in LD on chromosome 4q28.2. The x-axis is the base pair position and the y axis is the –log_10_ of the p-value for the association with Braak stage. The blue line represents the recombination rate. (**c**) Association between single-nucleotide polymorphism (SNP), rs56405341 and Braak tangle stage (adjusted for age and sex). Pairwise comparisons using Wilcoxon rank sum test, AA-AG *p*=0.024, AA-GG *p*=3.3E-05, AG-GG *p*=7.2E-05.

**Table 2.** Summary of SNPs reaching genome-wide suggestive and significance

| Chr | Base pair | SNP | Genes | A1 | Beta | SE | L95 | U95 | t-statistic | p |
| --- | --- | --- | --- | --- | --- | --- | --- | --- | --- | --- |
| 2 | 11935322 | rs78580932 | GREB1, NTSR2, LPIN1, MIR4262 | C | -0.54 | 0.10 | -0.76 | -0.34 | -5.15 | 3.55E-07 |
| 2 | 235768810 | rs74600760 | ARL4C, SH3BP4 | T | -0.53 | 0.11 | -0.74 | -0.32 | -4.86 | 1.48E-06 |
| 3 | 61518166 | rs111610564 | FHIT, PTPRG | C | -0.49 | 0.10 | -0.70 | -0.30 | -4.82 | 1.81E-06 |
| 3 | 140216029 | rs349509 | CLSTN2, CLSTN2-AS1, TRIM42 | A | -0.77 | 0.16 | -1.08 | -0.47 | -4.97 | 8.64E-07 |
| 3 | 179019705 | rs13323081 | PIK3CA, KCNMB3, ZNF639, MFN1 | G | 0.24 | 0.05 | 0.15 | 0.33 | 5.17 | 3.16E-07 |
| 4 | 130085480 | rs56405341 | C4orf33, SCLT1, JADE1 | A | 0.25 | 0.05 | 0.16 | 0.35 | 5.52 | **4.82E-08** |
| 4 | 137329297 | rs77506227 | - | A | -0.79 | 0.16 | -1.12 | -0.48 | -4.88 | 1.34E-06 |
| 5 | 150473104 | rs6579838 | GPX3, TNIP1, ANXA6 | C | 0.23 | 0.05 | 0.14 | 0.32 | 5.10 | 4.57E-07 |
| 10 | 124487608 | rs150945906 | DMBT1, C10orf120, DMBT1P1 | A | -0.64 | 0.14 | -0.91 | -0.38 | -4.77 | 2.31E-06 |
| 15 | 24515682 | rs147462127 | PWRN2, PWRN1 | T | -0.84 | 0.17 | -1.18 | -0.51 | -4.98 | 8.37E-07 |
| 15 | 36865219 | rs185845364 | C15orf41, CSNK1A1P1 | G | -0.79 | 0.16 | -1.11 | -0.48 | -4.94 | 9.79E-07 |
| 17 | 1879293 | rs12946465 | RPA1, RTN4RL1, DPH1, OVCA2 | C | -0.21 | 0.05 | -0.31 | -0.13 | -4.75 | 2.51E-06 |
| 19 | 45719790 | rs10415392 | TRAPPC6A, BLOC1S3, EXO3L2, MARK4 | T | 0.34 | 0.07 | 0.21 | 0.48 | 4.85 | 1.59E-06 |
| 20 | 56754110 | rs78923929 | C20orf85, PPP4R1L, RAB22A | A | -0.65 | 0.13 | -0.92 | -0.40 | -4.95 | 9.76E-07 |

genome wide significant difference in bold, chr=chromosome. SNP=single nucleotide polymorphism, SE=standard error

Replication of our SNP in an independent cohort proved challenging given the lack of datasets containing similarly neuropathologically ascertained individuals with PART. Nevertheless, we identified two other SNPs (rs4975209 and rs10009321) in the 4q28.2 locus in a prior Alzheimer disease (AD) GWAS that also used Braak as a quantitative trait, however they were not in high D’ with rs56405341, our lead SNP ^52^. We also observed in a separate AD GWAS using the cerebral spinal fluid Aβ42/40 ratio (dichotomized normal and abnormal) another SNP in the region (rs13129839) in high D’ with our lead and supporting SNPs (>0.89), at a genome-wide suggestive significance level and with a positive (protective) odds ratio (*p* = 9.0 × 10^−6^, OR = 0.043) ^53^. Taken together, these two independent genetic studies in AD suggest the signal in our GWAS might not be spurious.

We also examined candidate SNPs previously found to be associated with AD and progressive supranuclear palsy (PSP) in prior GWAS studies to explore convergent and divergent genetic risk (Table 3, Supplementary table 5). Of the 52 candidates investigated, we found five associated with PART (*SLC24A4, MS4A6A, HS3ST1, MAPT*, and *EIF2AK3*). rs12590654, which is associated with *SLC24A4*, had the highest significance level (*p*=0.001). rs1582763, rs2081545, rs7935829, were all associated with *MS4A6A* (*p*=0.01, 0.01, 0.02 respectively). The remaining AD SNP, rs7657553, was associated with *HS3ST1* (*p*=0.02). We found two variants that overlapped with PSP risk. rs242557 (*p*=0.02) in the *MAPT* locus, and rs7571971 (*p*=0.03) in the *EIF2AK3* locus. In summary, seven of the 52 probed risk AD and PSP SNPs showed significant associations (*p*<0.05) in PART.

**Table 3.**
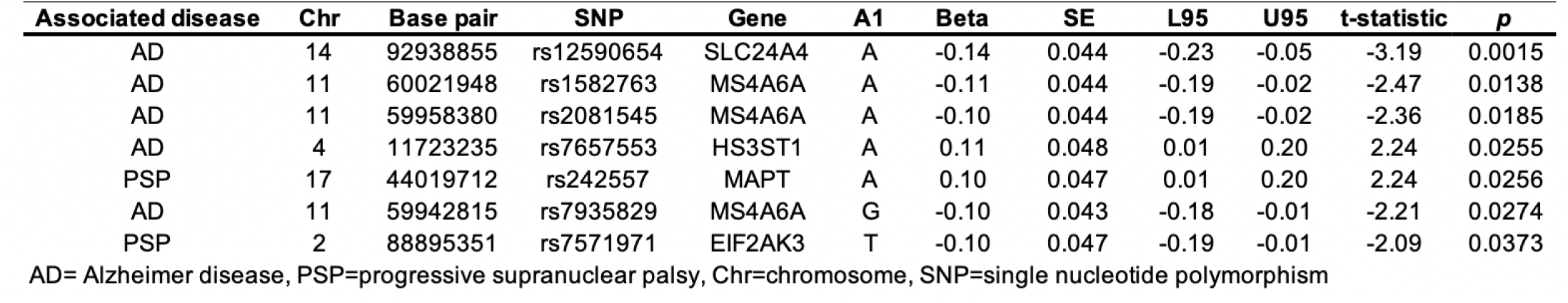
Significant overlapping PART genetic hits with Alzheimer disease and progressive supranuclear palcy

| Associated disease | Chr | Base pair | SNP | Gene | A1 | Beta | SE | L95 | U95 | t-statistic | p |
| --- | --- | --- | --- | --- | --- | --- | --- | --- | --- | --- | --- |
| AD | 14 | 92938855 | rs12590654 | SLC24A4 | A | -0.14 | 0.044 | -0.23 | -0.05 | -3.19 | 0.0015 |
| AD | 11 | 60021948 | rs1582763 | MS4A6A | A | -0.11 | 0.044 | -0.19 | -0.02 | -2.47 | 0.0138 |
| AD | 11 | 59958380 | rs2081545 | MS4A6A | A | -0.10 | 0.044 | -0.19 | -0.02 | -2.36 | 0.0185 |
| AD | 4 | 11723235 | rs7657553 | HS3ST1 | A | 0.11 | 0.048 | 0.01 | 0.20 | 2.24 | 0.0255 |
| PSP | 17 | 44019712 | rs242557 | MAPT | A | 0.10 | 0.047 | 0.01 | 0.20 | 2.24 | 0.0256 |
| AD | 11 | 59942815 | rs7935829 | MS4A6A | G | -0.10 | 0.043 | -0.18 | -0.01 | -2.21 | 0.0274 |
| PSP | 2 | 88895351 | rs7571971 | EIF2AK3 | T | -0.10 | 0.047 | -0.19 | -0.01 | -2.09 | 0.0373 |

AD= Alzheimer disease, PSP=progressive supranuclear palsy, Chr=chromosome, SNP=single nucleotide polymorphism

Next, we refocused on our strongest association at the 4q28.2 locus. Examination of RNA expression quantitative trait loci (eQTL) using the Brain-eMeta eQTL summary data (derived from six datasets; Genotype-Tissue Expression v6, the CommonMind Consortium, Religious Orders Study and Memory and Aging Project, the Brain eQTL Almanac project, the Architecture of Gene Expression, and eQTLGen) did not contain significant SNP associated eQTLs for any of the genes in the 4q28.2 locus ^54^. Because our GWAS quantitative trait was specific to tau pathology, we then examined mRNA expression levels of all the genes contained in the locus using a novel single-cell soma RNA sequencing dataset which measured transcriptomic changes specifically in tangle-bearing neurons and non-tangle-bearing neurons (Fig. 2a-c). Using this dataset, we found that tangle-bearing excitatory neurons significantly differentially expressed *JADE1* compared to non-tangle-bearing excitatory neurons (*p*=1.04×10^−61^). Conversely *C4orf33* and *SCLT1* had low levels of expression regardless of cell type and tangle status. Additionally, we observed an upregulation of *JADE1* expression in tangle-bearing inhibitory neurons however it was statistically underpowered due to low cell abundance and high variance. Furthermore, we observed two subpopulations of excitatory neurons in which *JADE1* was significantly differentially expressed (adjusted *p*=4.55×10^−15^, 7.82×10^−8^). We then examined the relative average expression and percentage of cells expressing *JADE1* and observed increases in both metrics in tangle-containing neuronal populations compared to non-tangle-containing neuronal populations (Fig. 2d-f). *SCLT1* and *C4orf33* had nominal expression levels in a substantially smaller percentage of neurons. Taken together, these data suggest increased *JADE1* expression is unique to specific populations of neurons which also contain NFTs. Because we did not observe significant differences in *C4orf33* or *SCLT1* expression across the two groups, and the signals were very low, these data suggest that *JADE1* as the strongest candidate gene in the locus.

**Fig. 2.**
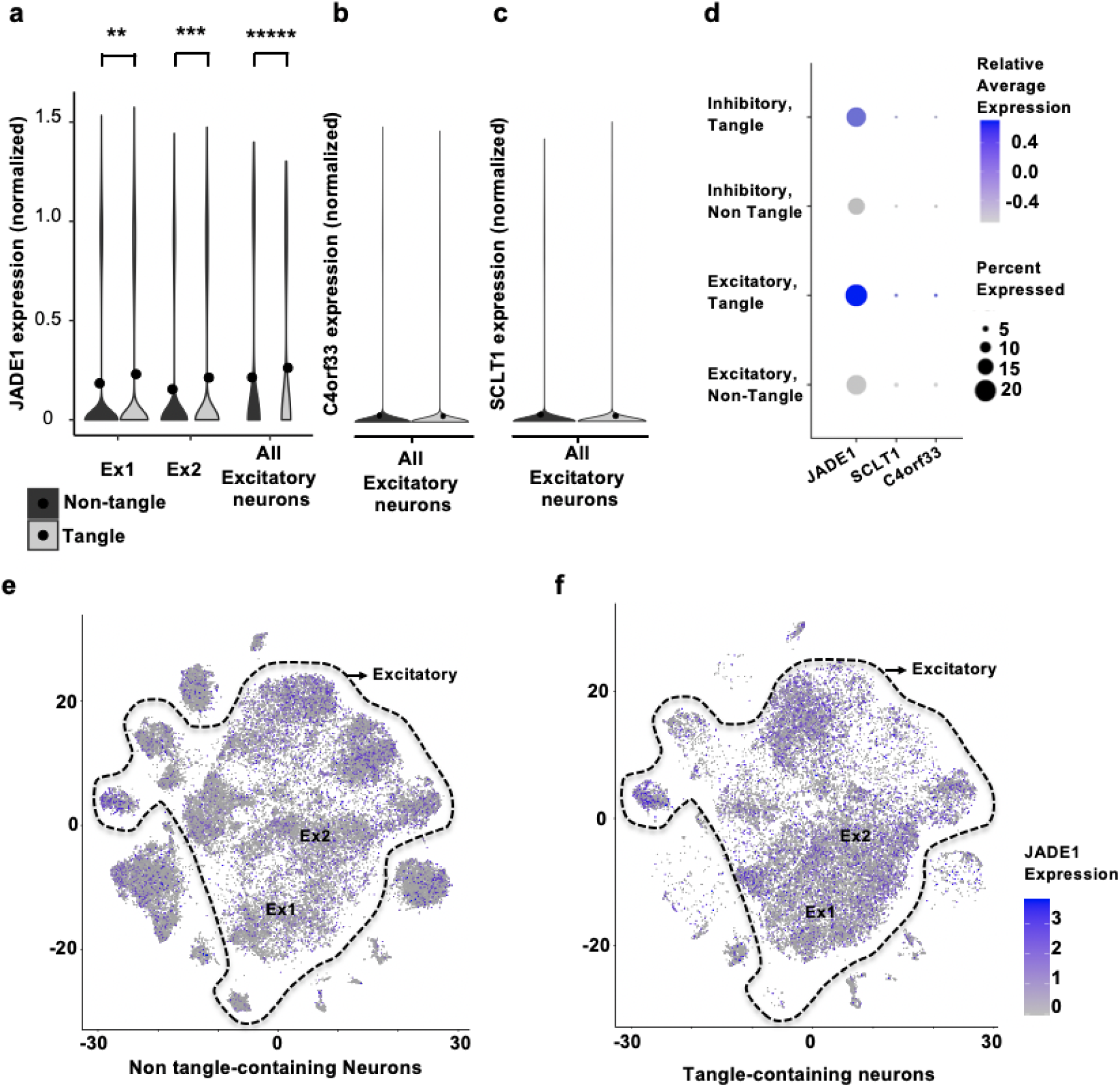
Single-cell sequencing of neurons with and without tau aggregates reveals *JADE1* mRNA is upregulated in tangle-bearing neurons. Neurons with and without neurofibrillary tangles from human post-mortem brains samples were separated using fluorescence-activated cell sorting and single-cell RNA-sequencing was performed. (**a**) In 2 unique excitatory neuronal populations *JADE1* mRNA was significantly differentially expressed in the tangle bearing neurons (adjusted *p*= 7.82×10^−8^, 4.55×10^−15^) and comparing the overall population of tangle-bearing excitatory neurons to non-tangle bearing neurons the value is highly significant (adjusted *p=*1.04×10^−61^). (**b**,**c**) The other two genes in the locus, *C4orf33* and *SCLT1*, were overall nominally expressed in both excitatory neuronal groups, as well as subclusters (data not shown). (**d**) A dot plot showing average relative expression and percent expression of the candidate genes in the locus. Both *JADE1* relative average expression and percentage of cells expressed was higher than *C4orf33* and *SCLT1*. (**e**,**f**) A t-distributed stochastic neighbor embedding (tSNE) plot showing the populations of neurons, tangle bearing status, and relative expression of *JADE1* in neuronal subpopulations.

Given our evidence that *JADE1* is genetically and transcriptionally associated with NFT pathology, we conducted an immunohistochemical study using specific antisera to JADE1 in our collection of post-mortem tauopathy brain tissue (Fig. 3). We assessed tauopathies that are known to involve preferentially tau isoforms with 3 microtubule-binding domain repeats (3R), 4 microtubule-binding domain repeats (4R) or a mixture of the two. We found strong and specific JADE1 immunopositivity in structures morphologically indicative of mature aggregate containing intracellular NFT in not only PART, but also the other mixed 3R/4R tauopathies (i.e., AD and chronic traumatic encephalopathy, Fig 3a-f). NFT pathology in PSP, corticobasal degeneration (CBD), and argyrophilic grain disease (AGD) were also immunopositive for JADE1 (Fig. 3g-l). Notably, gliofibrillary pathology in these diseases (i.e., aging-related tau astrogliopathy, tufted astrocytes, and astroglial plaques) were also immunopositive. Surprisingly, no signal was detected in NFT in Pick disease (PiD), a predominately 3R tauopathy (Fig. 3m,n). Double labeling experiments showed that early pre-NFT were negative for JADE1 suggesting that this factor begins to coalesce into NFT at the transition to the aggregate stage (Fig. 3o). To ensure the antibody was specifically targeting the JADE1 peptide and not binding non-specifically to NFT pathology, we blocked the JADE1 antibody and did not observe any staining in neurons that had the morphological features of an NFT (Fig. 3p). These findings indicate that JADE1 protein expression is localized specifically to cells with mature NFTs. Furthermore, JADE1 upregulation does not occur in PiD, the only 3R tauopathy examined, suggesting isoform specificity of expression.

**Fig. 3.**
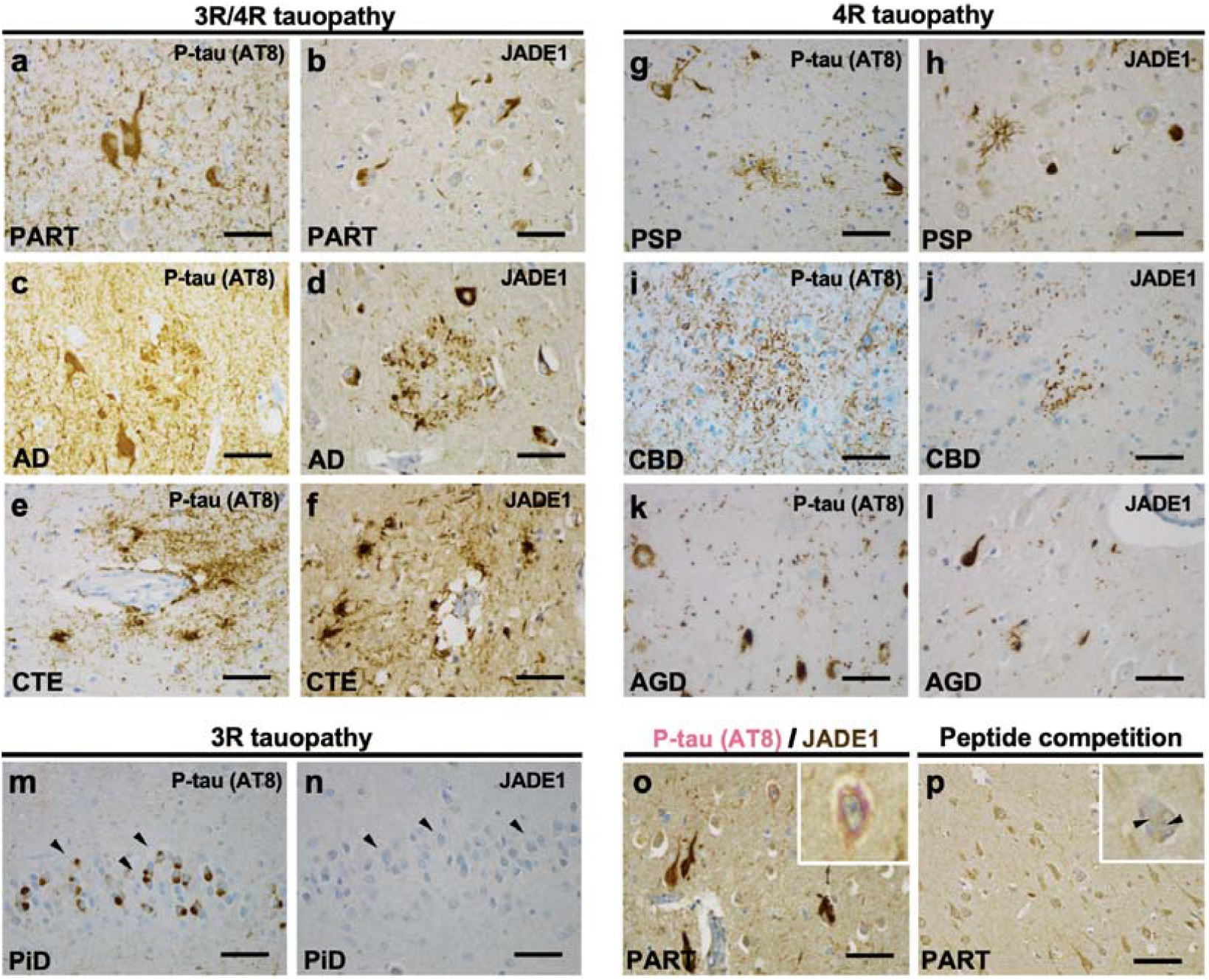
Selective immunolabeling of tau aggregates containing tau with four microtubule-binding domain repeats (4R), but not three (3R), in post-mortem human tauopathy brains with antisera targeting JADE1 protein. Immunohistochemical staining with phospho-tau (p-tau) specific antisera (AT8) and JADE1 specific antisera demonstrates neurofibrillary tangles (NFT) formation marked by the presence of JADE1 is specific populations of neurons and glia. (**a**,**b**) Primary age-related tauopathy (PART, *n*=3) NFTs contain JADE1 positive staining in the soma and neurites in the entorhinal cortex. (**c**,**d**) Alzheimer disease (AD, *n*=3) individual with beta-amyloid AT8-positive neuritic plaques and NFTs in the subiculum also display JADE1 immunopositivity in dystrophic neurites and NFTs. (**e**,**f**) Chronic traumatic encephalopathy (CTE, *n*=3) contains positive p-tau staining around a blood vessel in the depth of a neocortical sulcus that is also immunopositive for JADE1. (**g**,**h**) AT8 positive tufted astrocytes, oligodendroglial coiled bodies, and NFTs are positive in the subthalamic nucleus in a individual with progressive supranuclear palsy (PSP, *n*=3) which are also in immunopositive for JADE1. (**i**,**j**) Astrocytic plaques in corticobasal degeneration (CBD, *n*=3) and extensive thread-like pathology positive for p-tau and JADE1 in the neocortex. (**k**,**l**) In the cornu ammonis 1 (CA1) sector in a individual with argyrophilic grain disease (AGD, *n*=3), abundant grains that are immunopositive for p-tau and JADE1 are evident. (**m**,**n**) Pick disease (PiD), a 3R tauopathy, with Pick bodies in the dentate gyrus that are immunopositive for p-tau but negative for JADE1. (**o)** Double staining of a PART entorhinal cortex showing the absence of JADE1 (brown) staining in early pre-tangles, but the presence of p-tau (pink, see inset). (**p**) peptide competition demonstrating the antisera for JADE1 is specifically binding to the correct epitope give then absence of positive staining, but the presence of a tangle (inset). Scale bar 100 µm.

We then biochemically examined protein expression of JADE1 by western blotting using crude lysates derived from the entorhinal cortex and hippocampus proper (cornu ammonis 1-4 and dentate gyrus) of PART and AD individuals. JADE1 exists as 2 isoforms, JADE1 long (JADE1L) and JADE1 short (JADE1S), both of which contain proline, glutamic acid, serine, threonine (PEST) domains and 2 PHD fingers. However, the long form is 333 amino acid sequences greater in length and contains an additional PEST domain as well as a nuclear localization signal (Fig 4a). A Western blot analysis revealed a strong signal in both brain regions and diseases for JADE1S, but no bands were observed in the expected weight for JADE1L, indicating the observed signal is specific to the short isoform of JADE1 (Fig. 4b). This signal is in agreement with the immunohistochemical data given we did not observe a nuclear JADE1 signal which would suggest JADE1L, the form containing the nuclear localization signal, was also expressed. Furthermore, cytoplasmic colocalization of JADE1 and NFTs immunohistochemically raises a possibility that they form a functional complex in tauopathy brains. To examine this, co-immunoprecipitation (IP) was performed using crude brain lysate from PART individuals as the input. We first IPed using JADE1 antisera and observed a banding pattern that suggests JADE1 pulls down tau near the molecular weight of the 0N4R isoform (Fig. 4 c). To confirm these results, we reverse co-IPed JADE1 with 0N tau antisera and observed a banding pattern indicating the JADE1S isoform was pulled down (Fig. 4d). To investigate the observed JADE1 immunoprecipitated lower banding pattern (Fig. 4c), we treated this form with protein phosphatase over time and observed a shift over time to the expected 58 kDa weight (Fig. 4e), indicating JADE1 anti-sera was not able to recognize the more abundant native phosphorylated JADE1S and instead a low abundant species of the protein. This data suggests the predominant form of the JADE1S input is likely being biologically modified (i.e., phosphorylation, acetylation, etc.). Staining with C-terminal isoform-specific anti-tau antibodies (3R and 4R tau) revealed that the co-immunoprecipitated tau was predominantly 4R, thus 0N4R tau (Fig. 4f,g). We then ran western blots for the IPed JADE1 using multiple phospho-tau site-specific antibodies including RZ3 (pThr231), PHF1 (pThr181), CP13 (Ser202), AT8 (Ser202, pThr205), S214 (Ser214) and PG5 (Ser409) of which multiple phosphor tau epitopes showed prominent reactivity (Fig. 4h). Lastly, to confirm protein-protein interactions of Jade1S and 0N tau, proximity-ligation assays (PLA) were performed in fixed hippocampus from PART individuals. The PLA technique utilizes one pair of oligonucleotide-labeled antibodies that detects different epitopes of the two proteins in close proximity (maximum 30-40 nm apart). The prominent red fluorescence signal surrounding the nuclei of a neuron indicates the close proximity of these two proteins (Fig. 4i), whereas control samples lacking one of the two antisera did not show any signal (Supplementary Fig. 7e,f). In summary, the biochemical data strongly indicates a JADE1S, 0N4R tau immunocomplex.

**Fig. 4.**
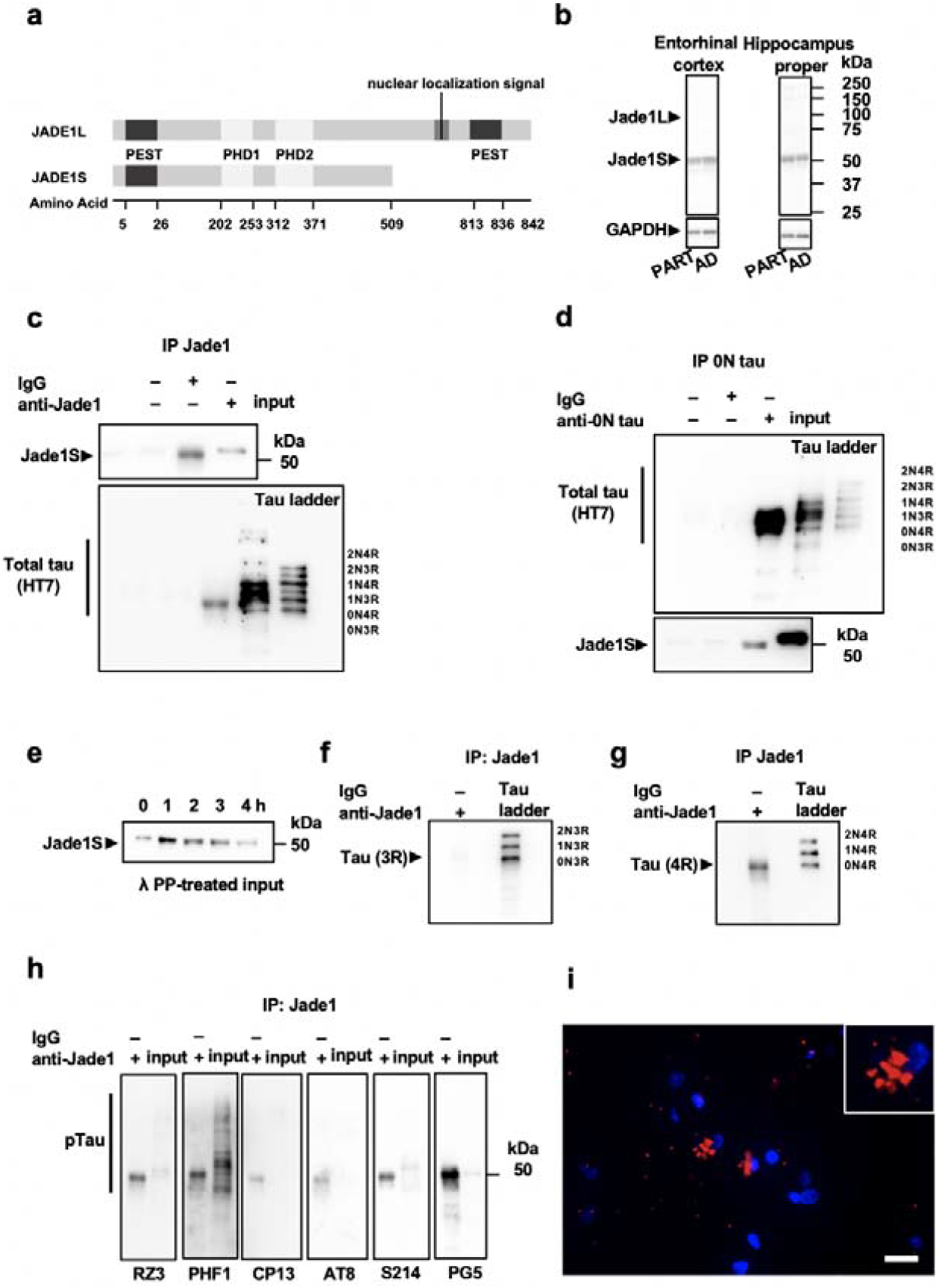
Biochemical analysis of JADE1S and total tau validates the interaction with 4 micr tubule-binding domain repeats (4R) but not 3R in post-mortem human brain tissue from tauopathy individuals. (**a**) Schematic of the two JADE1 isoforms, JADE1S and JADE1L (**b**) Representative immunoblot using antisera targeting JADE1 measured in entorhinal cortex and cornu ammonis in individuals with primary age-related tauopathy (PART) and Alzheimer disease (AD) shows a banding pattern with the JADE1S isoform at 58 kDa but not the JADE1L at 95 kDA. GAPDH was used as a loading standard. (**c**) Co-immunoprecipitation (co-IP) using JADE1 antisera pulls down tau near the molecular weight of the 0N4R isoform. (**d**) Reverse co-IP with 0N tau antisera pulls down the JADE1S isoform. (**e)** The pulled down form of Jade1S molecular weight shifts downward after treatment with lambda protein phosphatase over time to expected 58 kDa weight. (**f**,**g**) Co-IPed tau with JADE1 stained with C-terminal isoform specific anti-tau antisera are the 0N4R isoform and not the 3R isoform (**h**) Co-immunoprecipitated tau was positively stained with multiple phospho-tau specific antibodies with the largest signal coming from pSer396 pSer404 (PHF1), pSer214 (S214), and pSer409 (PG5). (**i)** Duo-link assay showing positive luorescence signal (red) around the nucleus of neurons (blue) indicating the potential interaction between JADE1 and 0N terminus tau detected using the corresponding two primary antibodies in the soma (inset). Scale bar, 20 μm

Lastly, we asked whether JADE1 plays a functional role in tau-induced neurotoxicity. We used a *Drosophila* model that overexpresses human mutant 0N4R tau^55^ as well as RNAi-mediated reduction of rhinoceros (rno), the highest matched JADE1 human ortholog in *Drosophila*. We first blindly evaluated the fly eye phenotype using a semi-quantitative assessment of eye size, roughness, overall shape, and conical shape and saw a significant increase in eye severity between tau transgenic *Drosophila* and tau transgenic *Drosophila* with rno knockdown (Fig. 5a-e, *p*=8.7 ×10^−5^). We did not observe significant differences between rno^rnai^ and control group in the absence of transgenic tau. To directly quantify neurodegeneration in the *Drosophila* brain, we quantified TUNEL-positive cells throughout the *Drosophila* brain. We find that rno knockdown significantly enhances neurotoxicity in tau transgenic *Drosophila* but is not sufficient to induce neurotoxicity in control flies based on TUNEL staining (*p*=0.008, Fig. 5f-i). These data provide *in vivo* evidence that JADE1/rno loss plays a mechanistic role in promoting neurotoxicity in tauopathy, and suggest that proper functioning of JADE1/rno is protective

**Fig. 5.**
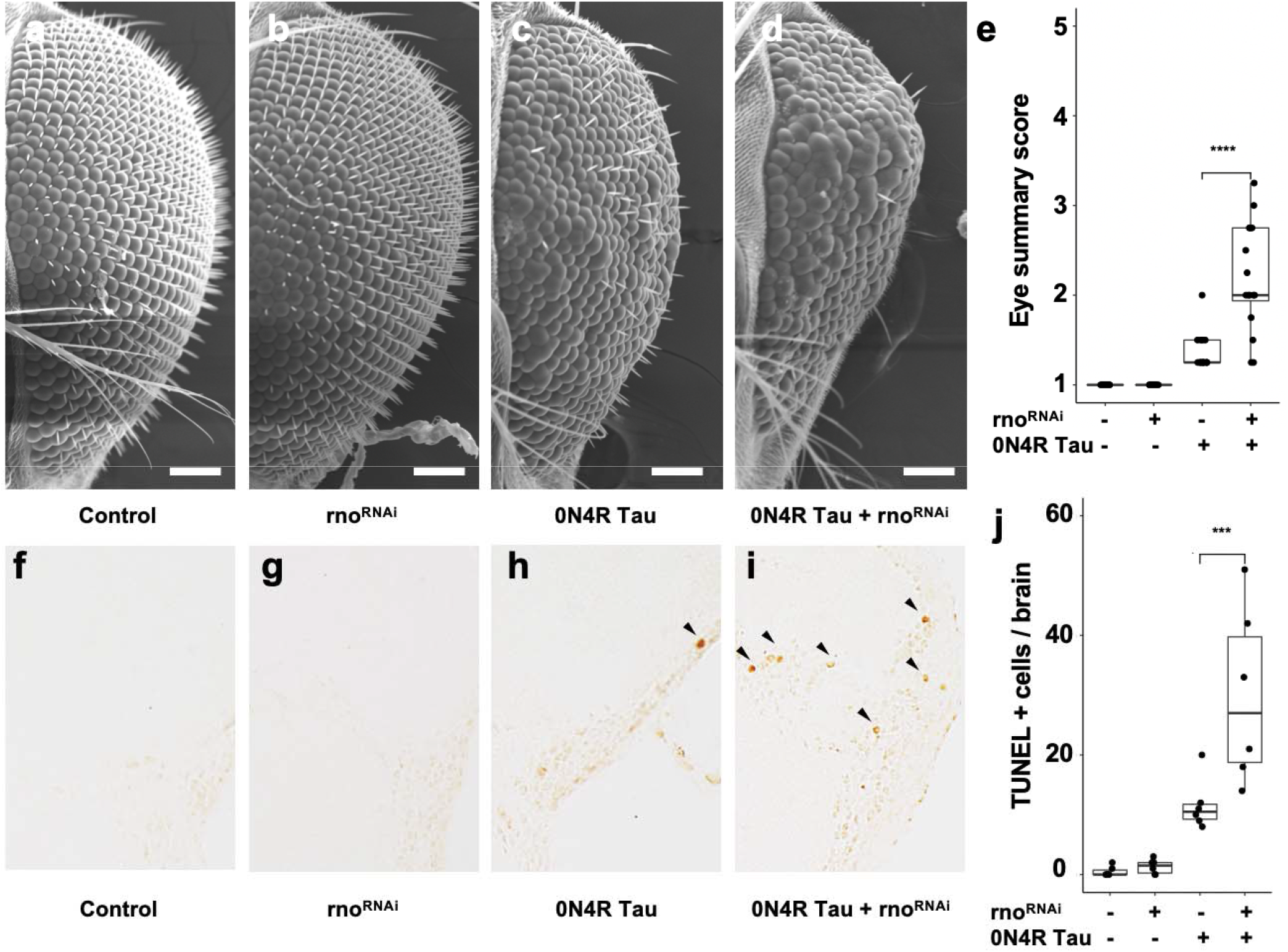
RNAi-mediated knockdown of rno enhances tau-induced rough eye and neurotoxicity in the adult *Drosophila* brain. (**a-e**) Rno^RNAi^ significantly enhances the tau-induced rough eye phenotype based on size, roughness, shape, and conical shape (*p*=8.7 ×10^−5^). (**f-j**) Rno^RNAi^ significantly enhances levels of terminal deoxynucleotidyl transferase dUTP nick end-labeling (TUNEL) in tau transgenic *Drosophila* compared to tau expressed alone (*p*=0.008). TUNEL was performed at day 10 of adulthood. An equal number of male and female flies were used for each experiment.

## Discussion

Genome-wide association studies (GWAS) have enabled advances in our understanding of sporadic tauopathies ^19,56-61^. Yet, in the context of the growing numbers of genes associated with Alzheimer disease (AD) ^32^, direct links with tau proteinopathy have been challenging to pinpoint as association signals show minimal overlap with factors classically implicated in tauopathy (e.g., proteostasis, tau protein kinases, etc.). This is not surprising given the ubiquity and heterogeneity of tau and other pathological changes in the aging human brain that are variably associated with cognitive impairment, which is the phenotype in most genetic studies despite the fact that it is a non-specific trait. We performed an autopsy-based GWAS, which minimizes classification errors and other issues with clinical studies of dementia, assembling the largest cohort of post-mortem brain tissues from aged individuals devoid of amyloid pathology with a goal of identifying factors independently associated with primary age-related tauopathy (PART). In doing so, we sought to provide genetic evidence that might clarify the controversial relationship between PART and AD, which it closely resembles neuropathologically. We failed to find an association with *APOE* ε4, the strongest common risk allele for sporadic late-onset AD. We did, however, find associations with other candidate loci in AD and progressive supranuclear palsy (*SLC24A4, MS4A6A, HS3ST1, MAPT* and *EIF2AK3*), as well as a novel genome-wide significant association at the chromosome 4q28.2 locus. Our data indicate that among genes in this locus, only the gene for apoptosis and differentiation in epithelia 1 (*JADE1*), a member of a small protein family that serves as a multifunctional adaptor implicated in renal and other cancers^62-64^, is upregulated in tangle-bearing neurons on both the mRNA and protein levels. This accumulation of JADE1 protein in NFT is not specific to PART but occurs in AD and all tauopathies with accumulation of 4R isoforms, but not in Pick disease which is a 3R tauopathy, indicating that this is generally a shared feature. We also show that JADE1 binds 0N4R tau, an isoform proposed to be a critical driver of tau pathology^65,66^. Finally, experiments in *Drosophila* show that reducing expression of the *JADE1* homolog rhinoceros (rno) exacerbates tau-induced neurotoxicity *in vivo*. Together, these findings strongly argue that JADE1 is a factor broadly capable of protecting neurons from neurofibrillary cell death that links PART to the tauopathic component of AD.

We confirm that the genetics of PART has a partial overlap with sporadic late-onset AD and replicated the consistent finding showing the lack of a signal in the *APOE* locus despite its strong association with AD ^32,67-70^. It has been shown previously that PART individuals have a higher *APOE* L2 allele frequency which distinguishes PART neuropathologically and genetically from AD ^71-73^. We have reported the frequency of the *APOE* ε4 allele is lower in PART^28^ and other studies have revealed similar findings in independent cohorts ^6,33^. It should be noted that these values can fail to reach significance when cross comparing groups with varying degrees of neuritic plaque pathology ^33,74^. Recent studies in mice and humans have indicated that the *APOE* ε4 allele may exacerbate tau pathology independently of Aβ plaques^75^, however other human studies failed to show an interaction^76,77^. These results add to the strong evidence that PART is entirely independent of *APOE* ε4 regardless of amyloid plaque pathology.

17q21.31 *MAPT* locus is the strongest genetic risk factor for PSP ^26^, which we and others had previously reported is also associated with PART ^28,31^. The *MAPT* H1 haplotype has also been associated with AD ^78-80^, however this region has a complex haplotype structure and may be more important in specific AD subgroups given the modest signal and variable findings in these association studies^81,82^. Intriguingly, in one AD GWAS using clinically ascertained individuals, removal of *APOE* ε4 carriers enhanced signals in the 17q21.31 locus ^23^. In the present study, there was only a modest association of *MAPT* with PART. This result may stem from differences in cohort selection with previous studies focusing on extremes, while we included a range of pathological severity, especially mildly affected individuals. Together, these data highlight that further investigation of the role of 17q21.31 *MAPT* locus in PART is warranted.

With regards to genes other than *APOE* and *MAPT*, we found four additional association signals in PART that overlap with either AD or PSP. Eukaryotic translation initiation factor 2 alpha kinase 3 (*EIF2AK3*) encodes an endoplasmic reticulum (ER) membrane protein critical for the unfolded protein response (UPR)^83,84^. Activation of the UPR has been observed and positively correlated with tau pathology, but not with Aβ plaque burden, in the hippocampus of aged cognitively normal individuals ^83^. Solute carrier family 24 member 4 (*SLC24A4*), a gene in the locus most strongly associated with PART and AD, is a member of the potassium-dependent sodium/calcium exchanger protein family and is involved in neural development, however little is known about its possible function in AD ^85,86^. Additionally, we identified an association of PART with the membrane spanning 4-domains A6A (*MS4A6A*) locus, which contains the binding regions for the transcription factor PU.1 which is selectively expressed in brain microglia and macrophages ^87^. The last overlapping genetic locus that contains heparan sulfate-glucosamine 3-sulfotransferase 1 (HS3ST1), has been suggested to modulate heparan sulfate proteoglycans as receptors for the spreading of tau ^88,89^. Taken together, these data are compatible with the hypothesis that the genes we identified in our GWAS are possible modulators of tau pathology.

Our study highlighted a novel locus on chromosome 4q28.2, that previously gave a suggestive signal in an independent autopsy GWAS in AD that also used Braak NFT stage as an endophenotype and a second GWAS focusing on dichotomized CSF Aβ positivity^53,90^. This prompted us to focus on this locus for further validation and functional studies. Because we failed to identify expression quantitative trait locus in both blood and brain datasets, we hypothesized that given our trait was tangle-specific, modulation of mRNA expression of genes in the locus might also be cell-type specific. This hypothesis was also motivated by the increase in genetic to transcriptomic associations found in cell specific populations in other contexts ^91-93^. Our results indicate that of the genes in the 4q28.2 locus, only *JADE1* mRNA was significantly and differently expressed in tangle-bearing neurons. Our immunohistochemical studies also showed JADE1 protein accumulation in both neuronal and glial tangle containing cells, validating these findings. Thus, JADE1 is most likely responsible for the GWAS signal at this locus.

Our immunohistochemical studies indicate that *JADE1* is potentially involved broadly in tauopathies. We observed immunopositivity not only in PART tangles, but also in tauopathies with aggregates containing 4R tau and mixed tauopathies with aggregates containing both 3R and 4R. The absence of staining in Pick disease, the only tauopathy with 3R tau aggregates examined, was surprising. Our biochemical studies suggest that JADE1 protein specifically interacts with 0N4R tau that is phosphorylated on epitopes known to be hyperphosphorylated in NFT. Our proximity ligation assay confirms the direct interaction between JADE1 protein and tau. Studies using cryo-EM and mass spectrometry have shown the ultrastructure of tau aggregates at unprecedented resolution, and it has been reported that 0N4R has a unique single β-sheet conformation for the fibril core ^94-96^. Intriguingly, recent mass spectrometry profiling studies of human postmortem brain tissues have suggested that changes in 0N4R tau isoform specifically is an early event in tauopathy ^97^. Double labeling experiments indicate that JADE1 increases shortly after the pre-tangle stage, accumulating alongside insoluble tau aggregates during the transition to the intercellular tangle stage perhaps reflecting a reactive/protective compensatory role ^98^.

Our findings provide important clues as to how *JADE1* may be functioning in tauopathy. *JADE1* has been previously implicated as a renal tumor suppressor involved in apoptosis, well as inhibition of Wnt signaling by ubiquitylating β-catenin, and interactions with cell-cycle regulators ^62,99-103^. Of the two JADE1 isoforms, only the short form which lacks a nuclear localization signal was observed, which was consistent with the cytoplasmic localization of the protein we observed by immunohistochemistry. This prompted us to hypothesize that JADE1S functions in the cytoplasm to mediate tauopathy. Our *in vivo* studies in which we reduced rno (the closest JADE1 ortholog) levels in *Drosophila* overexpressing mutant human 0N4R tau significantly enhanced the tau-induced rough eye phenotype as well as TUNEL-positive cells, a marker of apoptotic DNA fragmentation. While JADE1 has been shown to promote apoptosis in some contexts, our RNAi knockdown experiments suggest that proper functioning of JADE1 may be neuroprotective. Consistent with this, other studies have demonstrated that loss of rno function attenuates apoptosis in *Drosophila*^104^. Because previous studies have found JADE1 to be stabilized by the von Hippel-Lindau tumor suppressor which is a component of an E3 ubiquitin-protein ligase activity, JADE1 may be working with 0N4R tau through a similar mechanism to promote ubiquitin-mediated clearance of tau ^105-107^.

Our study has several notable limitations. Our sample size is small for GWAS standards; however, it should be noted that it is still the largest study of its kind. Given that PART is currently only reliably diagnosed postmortem, the study is limited by the availability of tissue that meet our strict criteria. Our study also relies on neuropathological assessments performed at multiple centers that may cause batch effects. While Braak staging is highly reproducible, with one report showing that across brains and raters the kappa score was greater than 0.90 ^108^, it is a semiquantitative (ordinal) variable. Further studies using a more quantitative approach to measuring tau burden that we have shown more closely align with functional clinical measures in PART may reveal additional candidates ^109^. Additional follow-up studies in experimental models are necessary further to validate our findings.

In conclusion, therapies for AD in clinical trials are moving towards targeting tau due to the lack of clinical efficacy of Aβ modulating therapeutic approaches ^110^. Here, by focusing on individuals with PART who lack Aβ plaques, we enriched our cohort for signals related to tau proteinopathy. Our analysis provides additional evidence that PART overlaps with but has considerable differences from AD. This interdisciplinary approach led to the identification of *JADE1*, which interacts with 0N4R tau, and is protective *in vivo*. Thus, JADE1 is a potential novel biomarker that differentiates tauopathies. Further understanding the genetics of PART will provide pathways for rationally designed therapeutics for degenerative tauopathies.

## Acknowledgments

The authors would like to acknowledge the neuropathology core of the Massachusetts Alzheimer Disease Research Center, the Biosample Management Repository at Genentech/Roche, the brain repository at UCI, Knight Alzheimer Disease Research Center Neuropathology Core at Washington University School of Medicine, the neurodegenerative disease brain bank at the University of California San Francisco, the Neuropathology Brain Bank & Research Core at Mount Sinai, and the following people: Ryan Cassidy Bohannan, Chad Caraway, Allison Beller, Kim Howard, Suresh Selvaraj, Ward Ortmann, Ping Shang, Jeff Harris, and Chan Foong,

